# Exploiting mechanisms for hierarchical branching structure of lung airway

**DOI:** 10.1101/2023.03.23.533381

**Authors:** Hisako Takigawa-Imamura, Katsumi Fumoto, Hiroaki Takesue, Takashi Miura

## Abstract

The lung airways exhibit distinct features with long, wide proximal branches and short, thin distal branches, crucial for optimal respiratory function. In this study, we investigated the mechanism behind this hierarchical structure through experiments and modeling, focusing on the regulation of branch length and width during the pseudoglandular stage. To evaluate the response of mouse lung epithelium to fibroblast growth factor 10 (FGF10), we monitored the activity of extracellular signal-regulated kinase (ERK). ERK activity exhibited an increase dependent on the curvature of the epithelial tissue, which gradually decreased with the progression of development. We then constructed a computational model that incorporates curvature-dependent growth to predict its impact on branch formation. It was demonstrated that branch length is determined by the curvature dependence of growth. Next, in exploring branch width regulation, we considered the effect of apical contraction, a mechanism we had previously proposed to be regulated by Wnt signaling. Analysis of a mathematical model representing apical constriction showed that branch width is determined by cell shape. Finally, we constructed an integrated computational model that includes curvature-dependent growth and cell shape controls, confirming their coordination in regulating branch formation. This study proposed that changes in the autonomous property of the epithelium may be responsible for the progressive branch morphology.

## Introduction

Mammalian lung airways exhibit an intricate tree-like structure that contributes to the efficiency of respiratory function. The lung airways are formed by repetitive elongation and bifurcation of the epithelial tube, which is common in organs such as blood vessels, kidneys, and glandular organs. The architecture of major airways (such as the main bronchi, lobar bronchi, and segmental bronchi) shows a stereotyped pattern in humans [1–3], suggesting the existence of a mechanism to program the large-scale design. In mouse lungs, early-formed proximal branch segments are thick and long; in contrast, distal branch segments become thinner and shorter [4–6]. Unlike the botanical tree, in which new branches are always thin, the lumen size of the lung epithelial tip is large in the early stage and decreases as bifurcation proceeds. In addition, branching events occur more frequently as development progresses [4,7–9], suggesting that branch formation is controlled with the developmental phase to configure the hierarchical structure. Several biological and theoretical investigations have explored the principle of epithelial branching through epithelial-mesenchymal interaction [10–16], but little is known about how the macroscopic configuration is ordered. In this study, we aimed to address the regulation of changes in the branch diameter and segment length during lung development. Epithelial-mesenchymal interaction is mediated by diffusible molecules such as fibroblast growth factor (FGF), bone morphogenic protein (BMP), sonic hedgehog (Shh), and Wnt [17,18]. Signaling molecules control the spatiotemporal coordination of cellular proliferation, migration, differentiation, and shape, leading to the arrangement of various cells in the prescribed shape for tissue formation. During lung development, epithelial growth is mainly promoted by FGF10, which is expressed in the outermost boundary of the mesenchyme [11,19,20]. FGF10 also induces epithelial expression of Shh, which diffuses into the mesenchyme and adversely represses FGF10 expression [21,22]. Theoretical studies have shown that this feedback interaction leads to interfacial instability in the growing epithelium and promotes epithelial branching [23]. This is similar to the mechanism by which bacterial colonies grow under nutrient-deprived conditions, where regions of high curvature are exposed to residual nutrients and elongate faster than regions of low curvature [24,25]. The tendency of the tip to grow faster is a common mechanism for spontaneous branch formation, such as crystal growth and viscous fingering [26–28]. Branching and bud formation with patterned proliferative activity at branch tips have also been observed for epithelial explants in extracellular matrix gels with an unbiased FGF distribution [10]. Drawing from these experimental insights, the diffusion-limited growth of epithelium has been described using reaction-diffusion [13,29] or phase-field schemes [12,30]. The mechanism of how heterogeneity of growth activity emerges is unknown; thus, the profile of FGF signaling activity in the lung epithelium requires confirmation to verify the common concept of branch formation models.

Wnt is also crucial for lung development [18]. The Wnt/ β-catenin pathway mediates cell fate determination in the embryonic state. In the lung mesenchyme, Wnt2 and Wnt2b are expressed as master regulators of proximal and distal patterning, which are distinguished by the epithelial expression of SOX2 and SOX9, respectively [31,32]. Loss of Wnt2/2b expression results in lung hypoplasia in the absence of small distal branches [33,34]. Double mutation of transcription factors Tbx2 and Tbx3, which act downstream of Shh and upstream of Wnt, respectively, also severely reduces the number of distal tips by inhibiting mesenchymal proliferation [35]. These findings indicate that Wnt may contribute to the generation of distal branches in the lungs. We previously reported that Wnt switches epithelial development from the branching phase (called the pseudoglandular stage) to the alveolus-shaping phase (canalicular stage), emphasizing the importance of Wnt signaling in distal duct morphology [36]. It was also revealed that activation of the Wnt/β-catenin pathway induces apical constriction of the lung epithelium and eventually scales down the bud size that emerges in mesenchyme-free epithelial cyst culture. Bud formation is an *in vitro* model of branch formation; therefore, our results indicate that Wnt signaling assists FGF10 in branching events. These results are consistent with a previous report showing that deletion of the Wnt receptor Fzd2 causes branching defects and cystic lung *in vivo* [37]. In this study, we considered how the interplay of growth control by FGF10 and cell shape regulation by Wnt can adjust the branch scale.

There are two distinct modes of lung branching: tip splitting and lateral branching [4]. The former is the symmetrical bifurcation at the distal end of a growing branch, and the latter is the sprouting in the middle of a branch. Stepwise observation of mouse lung development revealed that lateral branching mainly occurs during the early phase in the pseudoglandular stage, suggesting that lateral branching mainly generates asymmetric major branches. In contrast, tip splitting sequentially occurs in different sizes and frequencies during branch elongation, suggesting that the regulation of tip splitting is crucial for the hierarchical structure of branches. Accordingly, we focused on tip splitting in the pseudoglandular stage.

This study aimed to explore how branch formation is regulated in the lung through experimental and theoretical examinations. Experimental verification of the protrusion-grows-faster concept in the lung epithelium provides new insights into the tip-splitting branching frequency change across developmental stages through FGF. We also observed the change in cell size across the stages to explain the difference in tube size. Mathematical models demonstrate how these observed properties contribute to the hierarchical structure of the airways. Furthermore, we propose that apical constriction can efficiently promote tip splitting of thin branches to regulate their length based on observation of cell shape changes and mathematical analysis.

## Materials and Methods

### Animal

Transgenic mice expressing ERK FRET biosensors [38] were kindly provided by Michiyuki Matsuda (Kyoto University, Kyoto, Japan) through the JCRB Laboratory Animal Resource Bank of the National Institute of Biomedical Innovation. ERK FRET biosensors, EKAREV-nuclear export signals, are localized in the cytoplasm. ICR mice (Japan Charles River, Yokohama, Japan) were crossed with EKAREV mice, and the embryos were used for ERK activity measurement. ICR embryos were used for FGF10 uptake assay and immunostaining. All procedures involving animal experiments were reviewed and approved by the Ethics Committee of Animal Experiments, Kyushu University Graduate School of Medical Sciences (A25-182, A27-071, and A29-036).

### Mesenchyme-free culture of embryonic lung epithelium in Matrigel

Embryonic mouse lungs (E12.5, E13.5, and E14.5) were dissected and washed with Hank’s balanced salt solution (HBSS). Lung lobes were cut into pieces and soaked in dispase solution (1 U/mL in HBSS; Gibco) for 10 min at 37 °C. The tissues were washed twice in 10 mL of HBSS. Repetitive pipetting of tissues in HBSS broke up the mesenchyme and small epithelial explants remained. The supernatant containing mesenchymal cells was removed using a Cell Strainer with 40-µm pores (Falcon). The epithelial explants were picked using a micropipette under a stereomicroscope and placed on the bottom of a 35-mm glass-bottomed dish (IWAKI, blanched explants) or the µ-Slide Angiogenesis slide (*ibidi*, small explants for FGF10 dose-dependent assay). Matrigel (Corning, No. 356237) containing recombinant human FGF10 protein (PeproTech) was overlaid on explants and allowed to solidify for 15 min at 37°C, followed by the addition of the medium (2 mL for 35-mm dishes, and 50 µm for a well of the angiogenesis slide). The tissue culture medium was DMEM/Ham’s F-12 (Nacalai Tesque) supplied with Penicillin-Streptomycin Mixed Solution (×10, Nacalai Tesque), L-glutamine (584 mg/mL), and bovine serum albumin (10%). Samples were incubated for 18 h before ERK activity measurement. The FGF10 concentration is indicated as the mass concentration in the volume of the gel and medium.

### Quantification of ERK activity

We used a 405 nm laser for excitation and acquired the 475 nm and 530 nm emissions as CFP and YFP signals, respectively. The acquired confocal images were analyzed using Fiji software (National Institute for Health). Local ERK activity was estimated as the ratio of the YFP signal to the CFP signal in a small region of interest (ROI) with a diameter defined based on epithelium thickness (Fig 1 and S1). The ROI was limited in the regions where the signal of the epithelial cross section was acquired in the confocal image (123 ROIs for E12.5, 108 ROIs for E13.5, and 77 ROIs for E14.5, Fig 2 and S1). Mean command in Fiji was used to reduce the effect of spatial noise before quantifying fluorescence intensity, and signal ratios were corrected for background signal. Next, 2D tissue curvature was calculated from a series of ROI canters. Curvature was calculated from the coordinates of three ROIs four apart along adjacent data points to capture the smooth tissue curvature. The 3D curvature was not successful because of tissue damage caused by prolonged image acquisition times. The correlation with linear regression between ERK activity averaged over five neighboring points and tissue curvature was calculated by using *Mathematica* (Wolfram Research, Inc.).

**Fig 1.**
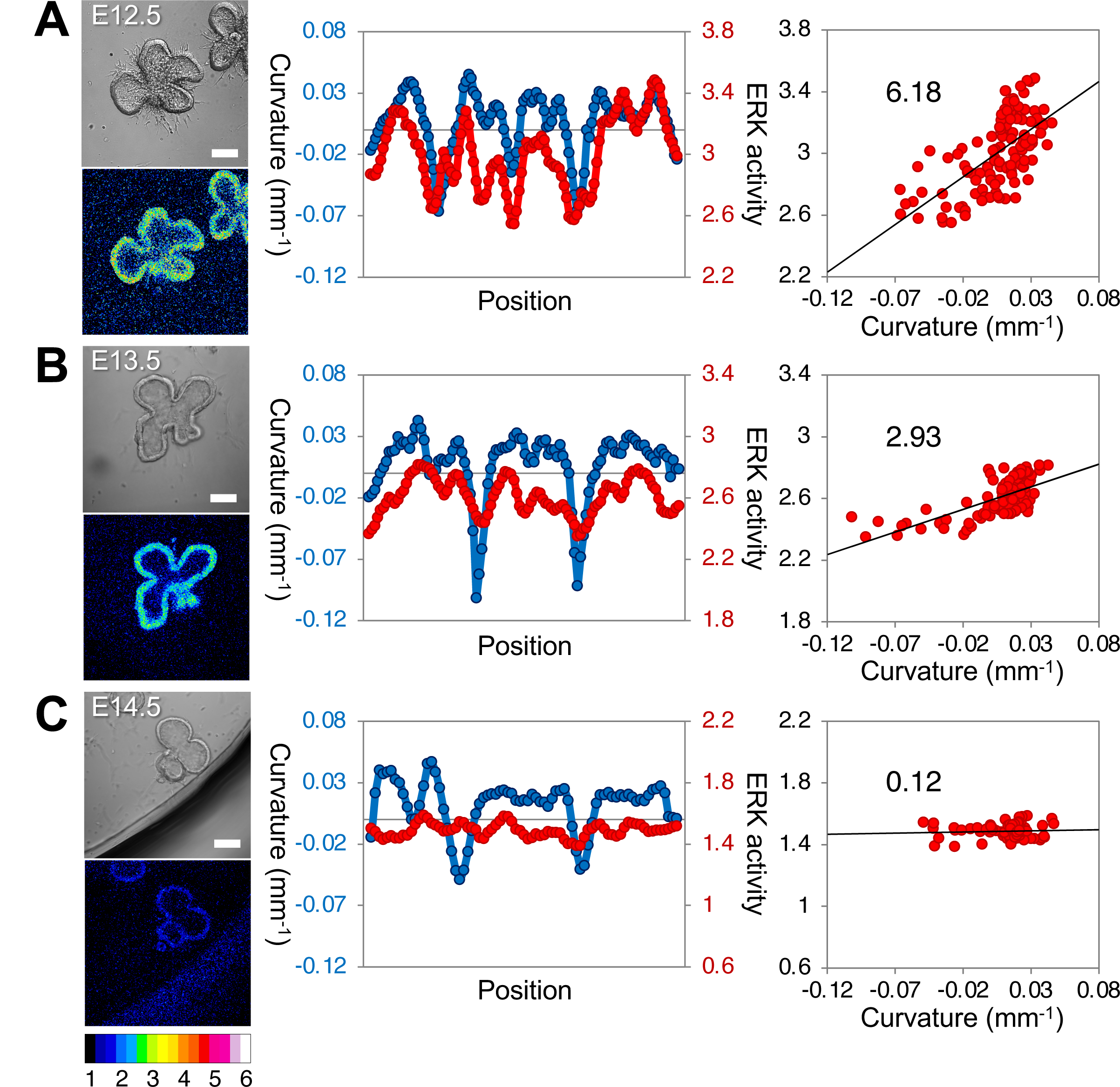
Map of ERK activity in embryonic mouse lungs. The upper left panel is the bright-field image, and the lower left panel shows the heat map representation of ERK activity. The middle panel shows cross-sectional tissue curvature (blue, left axis) and ERK activity (red, right axis) along the epithelial section. The right panel shows the correlation between the curvature and ERK activity. The color bar indicates the range of ERK activity. Epithelium explants at E12.5 **(A)**, E13.5 **(B)**, and E14.5 **(C)** were cultured with 500 ng/mL FGF10 for 18 h. The values in the figure represent the slope of the signal to curvature, determined through linear regression analysis (*R^2^*= 0.41 at E12.5, *R^2^* = 0.45 at E13.5, *R^2^* = 0.0031 at E14.5). Scale bars: 100 µm.

**Fig 2.**
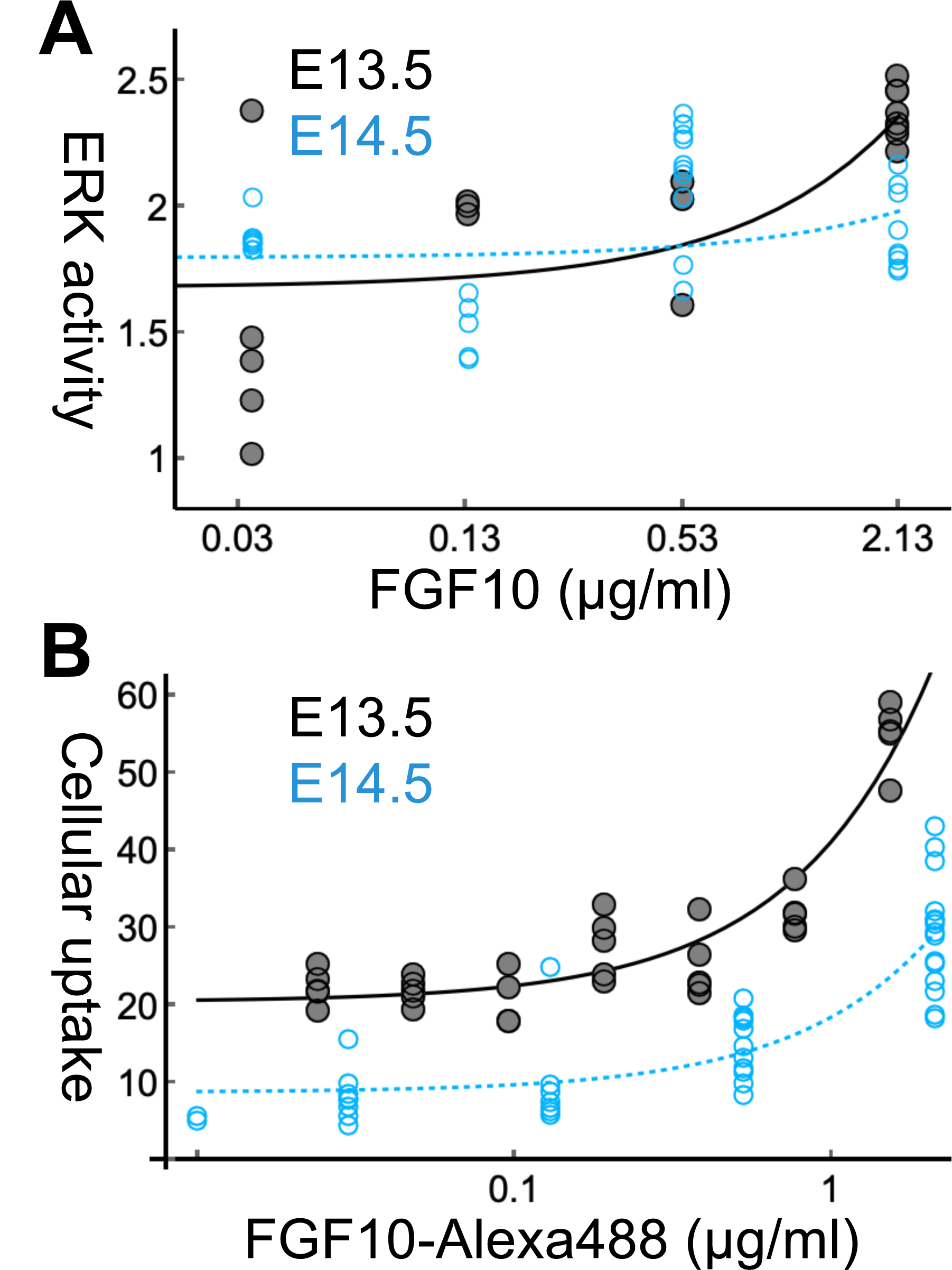
FGF10 dose dependency of the cellular response in E13.5 (black) and E14.5 (blue) epithelium. (**A**) The ERK activity of each cyst is represented by a dot in the semilog plot. The lung epithelium cysts of E13.5 and E14.5 (19 and 34 cysts, respectively) were cultured with FGF10 at concentrations of 0.03, 0.13, 0.53, and 2.13 µg/mL for 18 h. Linear regression analysis showed *y* = 0.330*x* + 1.68 with *R^2^* = 0.53 for E13.5 and *y* = 0.0334*x* + 1.89 with *R^2^* = 0.012 for E14.5. (**B**) The cellular uptake of FGF10-Alexa488 for 3 h was estimated for 34 and 44 explants of E13.5 and E14.5 lung, respectively. Each dot in the semilog plot represents a mean fluorescent intensity in an epithelial area of a cross-section. The concentration of FGF10-Alexa488 was 0.024, 0.048, 0.097, 0.19, 0.38, 0.78, and 1.55 for E13.5, and 0.03, 0.13, 0.53, and 2.13 µg/mL for E14.5. Linear regression analysis showed *y* = 20.7*x* + 20.4 with *R^2^* = 0.87 for E13.5, and *y* = 9.75*x* + 8.80 with *R^2^* = 0.72 for E14.5.

The ERK activity of the cyst (Fig 2A and panel A in Fig S2) was calculated as the ratio between the total intensities of YFP and CFP in the epithelial cyst area, excluding the luminal area in the cross-section. The FGF10 dose dependency was estimated by linear regression analysis, and its significant difference by developmental stage was tested using two-way analysis of variance (ANOVA) in the statistical software R (R Core Team).

### FGF10 incorporation assay

Recombinant FGF10 protein was labeled using an Alexa Fluor 488 Microscale Protein Labeling Kit (Invitrogen). To reduce the loss of physiological activity of FGF10, we optimized the labeling protocol and reduced the amount of Alexa Fluor used. The labeling reaction was conducted by mixing 100 µL of FGF10 solution (1 µg/mL in phosphate-buffered saline, PBS, pH 7.4), 5 µL of sodium bicarbonate (1 M), and 9.8 µL of Alexa488 solution (1:10 diluted) at 22°C. After a 15-min reaction, the unreacted dye in the sample was removed using a Slide-A-Lyser Mini Dialysis Unit (Thermo Scientific) in PBS for 16 h at 4°C. The purified solution was collected, and the protein concentration was measured using a NanoDrop spectrophotometer (Thermo Fisher). The FGF10-Alexa488 solution was stored in small aliquots at −20°C to avoid freeze-thaw cycles. Incubation and storage were performed in the dark.

Epithelial explants were embedded in Matrigel containing FGF10-Alexa488 protein in a 35-mm dish, which was subsequently filled with the medium. The protein incorporation was allowed for 3 h in a stage-top chamber (Tokai Hit) to maintain a 5% CO_2_-controlled atmosphere at 37°C followed by observation. The relative uptake activity was measured as the mean fluorescent intensity in the epithelial area of the cross-section of each explant. ROIs were defined as the round epithelial area, excluding the luminal area in the cross-section. While epithelial explants from E14.5 formed cysts within 3 hours, those from E13.5 exhibited slower cyst formation, with most regions of interest (ROIs) observed in sections from the tip regions (panel B in Fig S2).

FGF10 uptake was quantified as the mean intensity in the epithelial cyst area, excluding the luminal area in the cross-section. The FGF10 dose dependency was estimated by linear regression analysis, and its significant difference by developmental stage was tested using two-way ANOVA in the statistical software R.

### Immunostaining and quantification of epithelial thickness

Embryonic lungs from ICR mice were washed with PBS and fixed with methanol for 30 min. Permeabilization was in 0.1% Triton X-100 for 30 min, followed by the blocking with 1.5% normal goat serum for 60 min. The samples were stained with anti-mouse E-cadherin monoclonal antibody ECCD-2 (1:100, Takara) for 16 h at 4°C, followed by washing three times with 0.1% Triton X-100 in PBS. After incubation with Alexa Fluor 488-conjugated goat anti-rat IgG secondary antibody (1:200, Thermo Fisher) for 30 min, the sample was soaked in Sca*l*e reagent for tissue clearing overnight [39]. The cell nuclei were stained with 5 µg/mL propidium iodide (PI; Dojindo).

Thickness measurement was conducted by drawing small circles with diameters corresponding to the local thickness of the epithelium on the E-cadherin antibody-stained images. Circles were positioned to overlap halfway along the epithelium. The diameters of the circles were measured using Fiji and the mean and standard deviation were calculated to represent the epithelial thickness for each sample. The ROIs designated for thickness measurements are shown in Fig S3.

### Microscope observation

The lung explants were observed at 20× magnification using a Nikon C1 confocal microscope (Nikon Intech). Samples of live explants were kept in a stage-top chamber (Tokai Hit) to maintain a 5% CO_2_-controlled atmosphere at 37°C for cellular homeostasis during the observation.

### Particle cell model of the curvature-dependent growth

Particle-based two-dimensional (2D) model framework from our previous studies was inspired by particle-based methods for fluid simulation, incorporating interactions with Lennard-Jones-like potentials between particles [40–44]. In this study, the modeling framework was employed to describe epithelial growth and deformation. Each particle represents a cell (Fig 3). The cell-cell interaction and bending rigidity were introduced as previously described [42,43]. Curvature-dependent tissue growth was newly incorporated for this study and is mentioned in the Result section. Cell division event was described by replacing a cell with two daughter cells with a radius of 0.2, which gradually grew to the default mother cell radius of 0.4 in 1000 timesteps. Parameter values are in the supplementary table (S1 Table).

**Fig 3.**
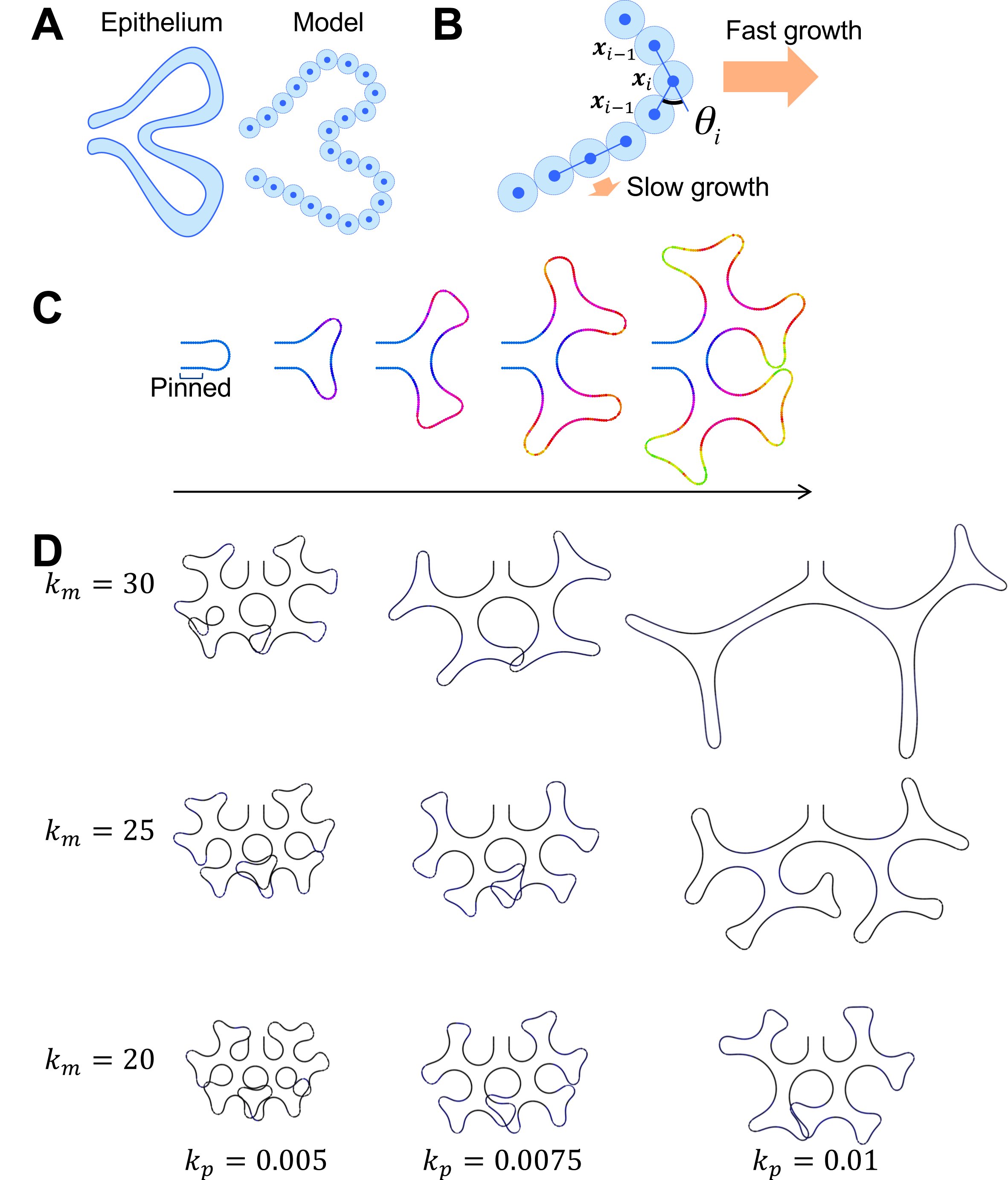
Protrusion growth described by the particle cell model. **(A)** The epithelial cell was represented by the particle in the model. The cell-cell distance is denoted by *d*. **(B)** Cell division was described by replacing a particle with two small particles, which gradually grow to the prescribed size. **(C)** The process of higher-order branching. Snapshots at *t* = 0, 75, 150, 225, and 300 are shown. The proximal end was pinned, and the distal tip was growing. The color indicates the cell generation. The number of cells was initially 50 and was increased to 450. **(D)** Branch scale control by curvature-dependent growth. Calculations corresponding to (C) were performed until the number of cells reached 800.

To obtain the initial geometry, 50 cells were arranged in a U-shape with a width of 5. The ten leftmost cells were pinned in their initial position. For relaxation of the shape of the cell sequence, cells were not grown, and the cell sequence was smoothed by bending rigidity until the shape became stationary. After sufficient relaxation time, numerical calculations of growth were performed. Pinned cells were not allowed to grow through the calculation, while growth was assumed for the remaining right tip side.

The time development of the model was calculated using C++ with Δ*t* = 0.005, and the graphics were generated using *Mathematica*. These computational procedures were used for other models throughout the study.

### Quadrilateral cell model with apical constriction

To describe tissue deformation coupled with cell shape control, we applied the model framework in our previous studies [36,45], where the cell shape is represented by four vertices (***a****_i_*, ***a****_i+_*_1_, ***b****_i_*, and ***b****_i+_*_1_ in Fig 4A) and the displacement of vertices are calculated. Three attributes are considered for cell shape: apical length, basal length, and cell area, each with its respective target value. A harmonic potential *W* and force ***f*** associated with the vertex **x** can be considered as follows:

**Fig 4.**
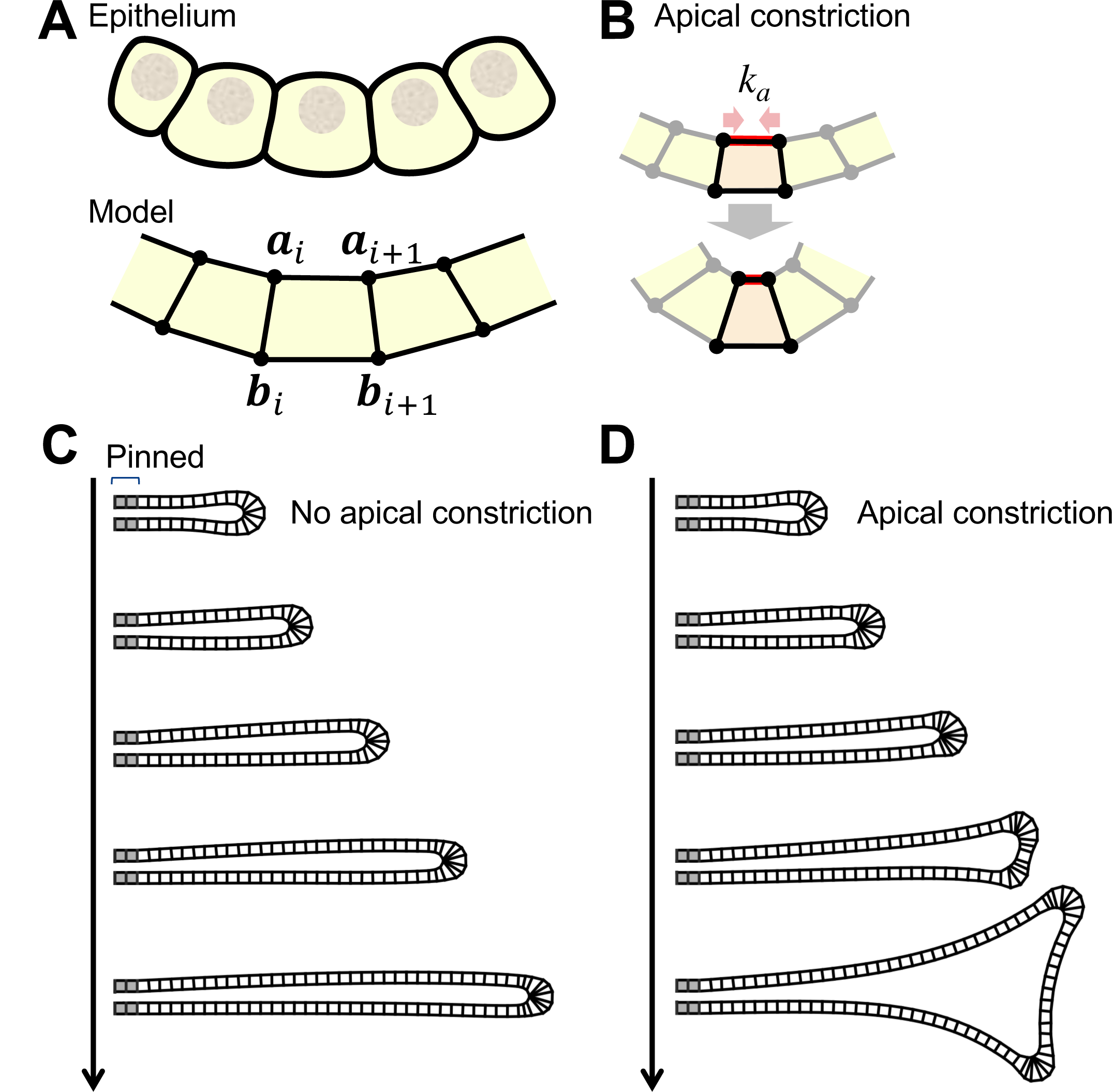
Effect of apical constriction on branch shape shown using the quadrilateral cell model. **(A)** The cross-section of epithelium is described as a sequence of quadrilaterals. The *i*-th cell was defined by four vertices ***a****_i_*, ***a****_i_*_+1_, ***b****_i_*, and ***b****_i_*_+1_. **(B)** Apical constriction is described by the attraction between apical vertices, with its strength represented by *k*_a_. **(C)** Curvature-dependent growth process. Snapshots at timesteps *t* = 0, 750 1750, 2750, and 3750 are shown. Pinned cells are indicated in gray. The case without apical constriction *k*_a_ = 0 is shown. **(D)** Apical constriction was introduced with *k*_a_ = 1.

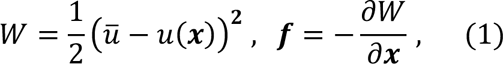

where *u*(***x***) is an attribute value and, *u̅* is its corresponding target value. The forces for each attribute were calculated by Eq (1). The target value of apical length is 0 for introducing apical constriction (Fig 4B). It was assumed that all cells exhibit uniform apical contraction for simplicity. The area regulation force based on isotropic restoration [36] tended to destabilize the cell shape, thus it was simplified to be based on adjusting the lateral length as follows:

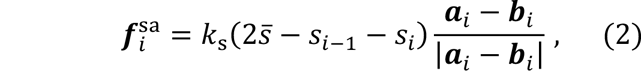

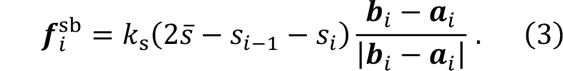

Eqs (2) and (3) are for the apical vertex ***a****_i_* and the basal vertex ***b****_i_*, respectively. The coefficient *k*_s_ determines the magnitude of the regulation, *s_i_* is the area formed by ***a****_i_*, ***a****_i_*_-1_, ***b****_i_*_-1_, and ***b****_i_*, and *s̅* is its target value. When the two cells sharing the vertex are symmetrical in shape, Eqs (2) and (3) are consistent with the energy-derived force [36].

To realize stable natural tissue shape, cell shape symmetry regulation was assumed for each cell, and bending rigidity was assumed for the sequence of the basal vertex as detailed previously [36]. Parameter values were adjusted for this study (S2 Table). We considered the displacement of the cell in the viscosity-dominated regime as follows:

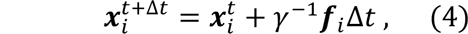

where ***x****_i_* is a non-dimensionalized position vector of the *i*-th cell center, *γ* is the friction coefficient, *t* is time, and Δ*t* = 0.005 is the timestep size.

Cell migration and division were assumed to depend on tissue curvature. Cell division was described by inserting new vertices into the middle of the edges of a fully grown cell at a probability *q_i_*. The daughter cells are set at half the size of the mother cells at the time of division and grow over 50,000 timesteps. Cell migration is described as the outward movement of vertices on the basal side represented by 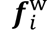. The dependency on the tissue curvature was assumed as follows:

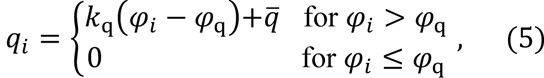

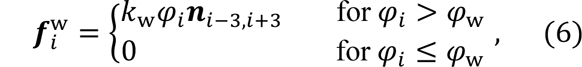

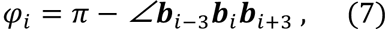

where *k*_w_ and *k*_q_ are the strength of curvature dependence, ***n****_i,j_* is the normal vector to ***x****_i_*-***x****_j_*, *φ*_w_ and *φ*_q_ are the critical angles for migration and cell division, and *q̅* is the basic probability of cell division. Tissue curvature was approximated using *φ_i_* as an alternative to *θ_i_*, aiming to avoid detecting local distortions.

The initial shape in Fig 4C and 4D was a thin tubular shape with an outer width of 3. The three leftmost vertices were pinned at the initial positions. After sufficient relaxation time, numerical calculations of growth were performed. Cells with pinned vertices were not allowed to grow through the calculation, while growth was assumed for the remaining right tip side.

In Fig 5A, epithelial cysts were depicted. No vertices were pinned, and no growth was assumed. Initially, *k*_a_ = 0 was set and increased by 0.003 at each timestep until *k*_a_ = 3.

**Fig 5.**
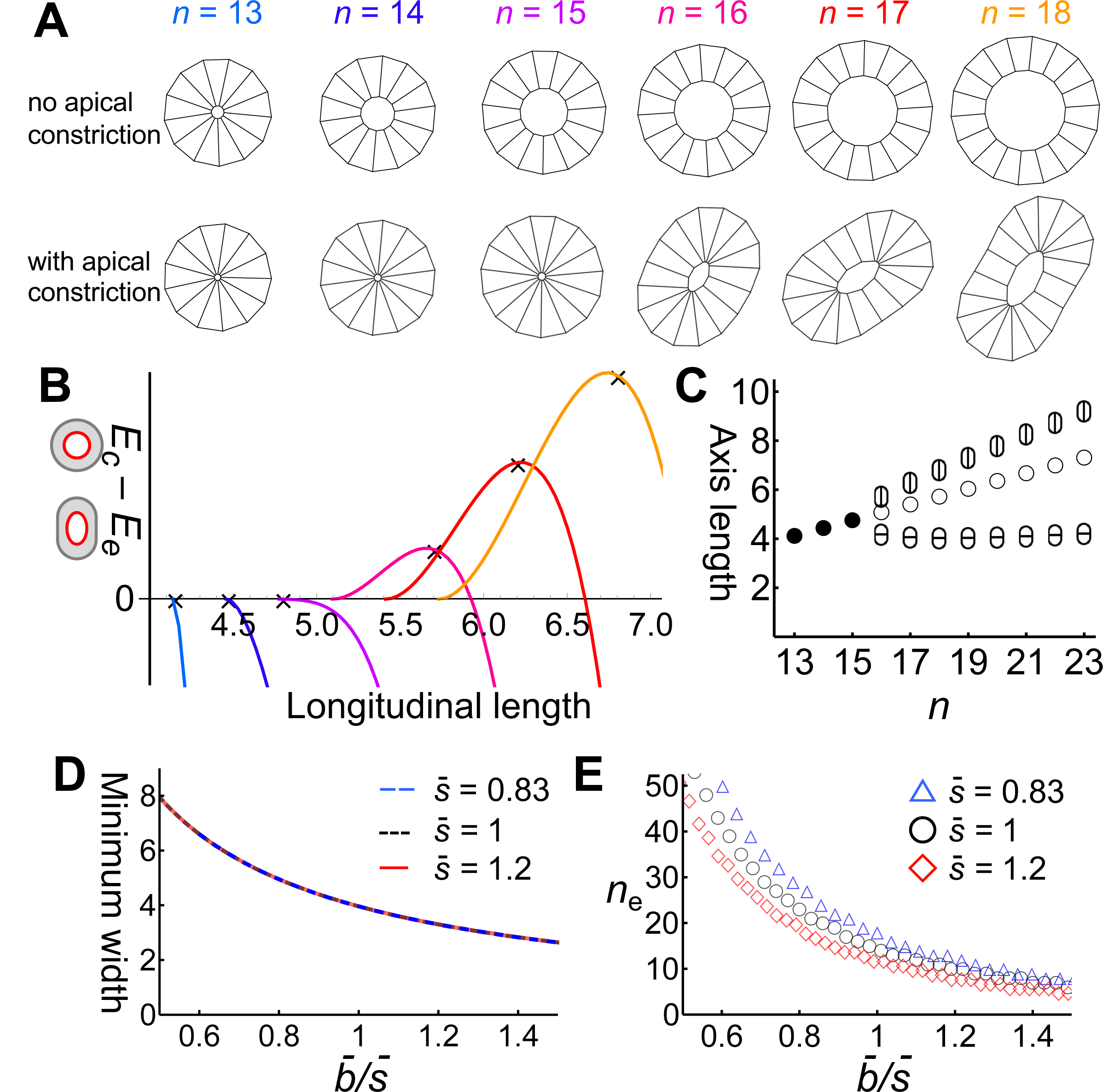
Mechanical instability of small cysts by apical constriction in the quadrilateral cell model. **(A)** Cyst of *n* cells was relaxed for a sufficient time (upper panels), then apical constriction was introduced and deformation was observed (lower panels). **(B)** The potential energy difference between the circular state (*E_c_*) and the elongated state (*E_e_*). Color corresponds to the cyst sizes in (A). The vertical axis is the longitudinal cyst length *L*. The minimum *L* corresponds to the diameter in the circular state. The cross indicates the cyst length under apical constriction in the numerical simulation (A), indicating that the energy calculation predicted the model behavior. **(C)** Estimated stable cyst shapes obtained from energy calculations plotted along the cell number *n*. When *n* ≤ 15, the stable shape was circular (closed circle). When *n* ≥ 16, the cyst was elongated, and the longitudinal length (ellipse with a vertical line) and the width (ellipse with a horizontal line) were plotted. The diameter of the corresponding circular cyst for *n* ≥ 16 was plotted for reference (open circle). **(D)** Estimated minimum cyst width for 13 ≤ *n* ≤ 27. The vertical axis represents J^’^/*s̅*, where J^’^ is the cellular basal length and *s̅* is the cellular area. Colors represent *s* values: 0.83 (blue dashed line), 1 (black dotted line), and 1.2 (red line). **(E)** Estimated critical cell number *n*_e_ for the shape transition from circular to elongated.

### Estimating mechanical instability in the cyst models

In the quadrilateral cell model, cysts were depicted in Fig 5A. Potential energy calculations were performed for these representations (Fig 5B–E), ranging from circular to elongated shapes, by considering the combination of a rectangle of length *x* and a semicircle of radius *y*. The longitudinal length of the cyst is *L* = *x* + 2*y*. It was assumed that the basal length and cell area are uniformly maintained as *b̅* and *s̅*, respectively. The sum of the basal lengths in the cyst is *b̅n* = 2*x* + 2*πy*. The apical lumen was approximated as an ellipse with a major radius *α* and a minor radius *β*. We estimated *α* and *β* by considering the conservation of cell areas and the relationship between cell height and apical length. The apical length of the cell with the highest curvature is denoted by *a_y_*. We assumed *a_y_*/*a* = *b̅*/(*x*/2 + *y*) by considering the cell shape as a trapezoid. The area of the cell is given by *s̅* = (*a_y_* + *b̅*)(*y* + *x*/2 − *α*)/2, where *α* was derived as in Eq (8) below. Additionally, given that the elongated cyst area is equal to the circular cyst area, expressed as *ns̅* = 2*xy* + *πy*^2^ − *παβ*, *β* was derived as in Eq (9). Eqs (8) and (9) are as follows:

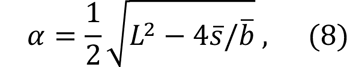

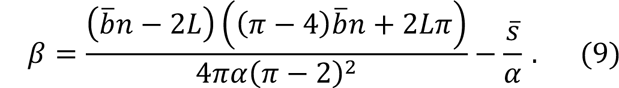

### Cell type switching model

The model dynamics of integrated (Fig 6) is based on the quadrilateral cell model, with parameter values provided in the supplementary table (Table S3). The cell is initially of the tip type and switches to the duct type when *φ_i_ > φ*_d_. The regulation of apical length and cell area were assigned for the tip type (*k*_at_, *a̅*_*t*_, and *s̅*_*t*_) and duct type (*k*_ad_, *a̅*_*t*_, and *s̅*_*t*_), respectively. The triangle-like shape of the tip type was described by setting the optimal apical length *a̅*_*t*_ = 0. The tall columnar shape of the duct type was characterized by setting *s̅*_*t*_ < *s̅*_*t*_ for the optimal cell area and equalizing apical and basal lengths as *a̅*_*t*_ = *b̅*. Migration was assumed only for the duct type and described by moving the basal vertex with force 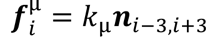. The difference in the epithelial thickness was described by changing *b̅*.

**Fig 6.**
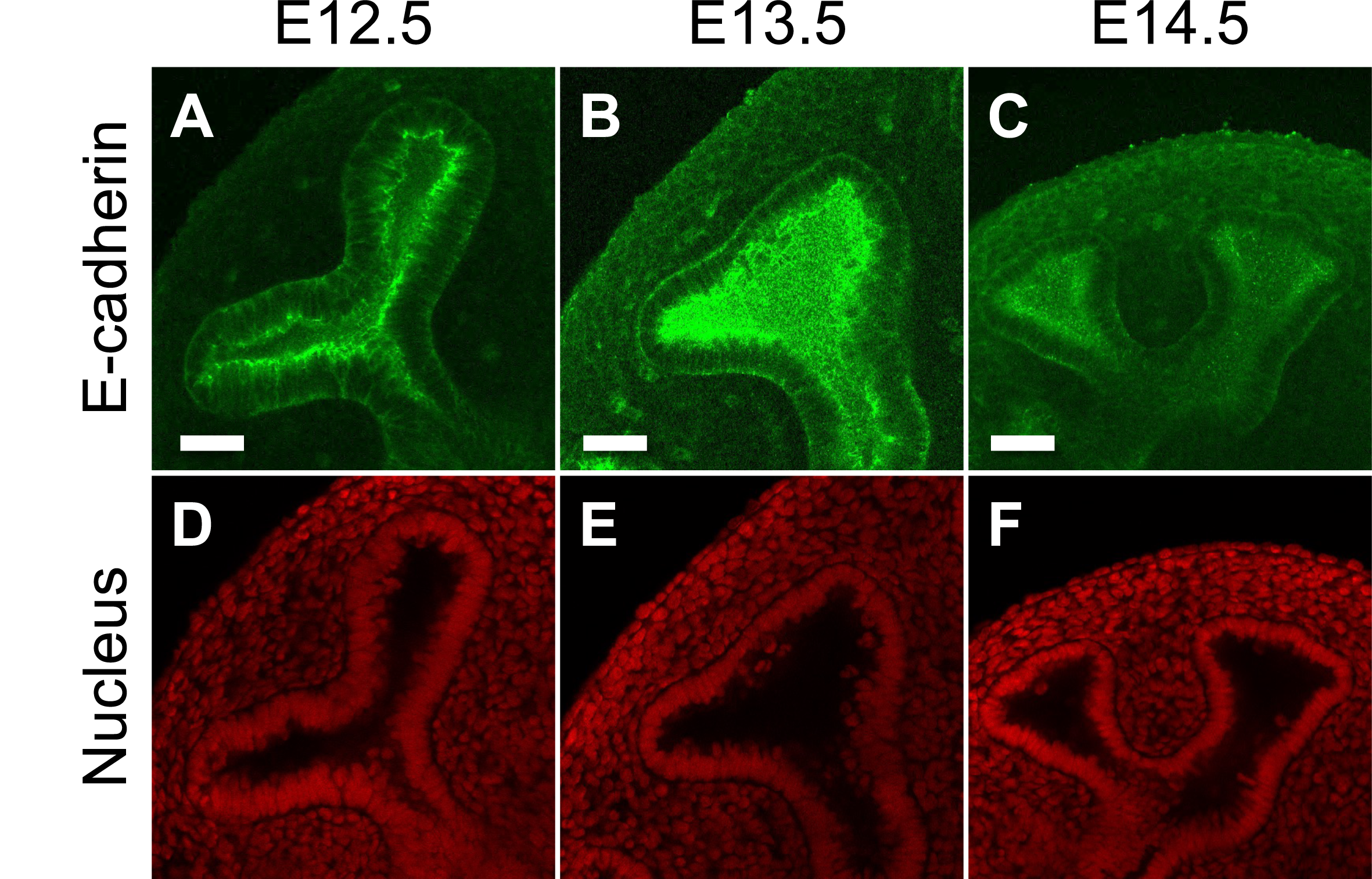
Changes in epithelial thickness during lung development. **(A–C)** E-cadherin staining was performed to show the epithelial shape of the growing tip at E12.5 (A), E13.5 (B), and E14.5 (C). **(D–F)** Corresponding nuclear staining with PI is presented. Scale bar: 100 µm.

The initial shape was assumed to be the tip of an epithelial branch with a width of 10, in which 13 cells at the tip were assumed to be the tip type and the others were assumed to be the duct type. Cells at the proximal ends were pinned at their initial positions and no cell division was assumed in these cells.

## Results

### Verification of the enhanced FGF10 signaling in protruded regions of lung epithelium

The assumption that the protruded region of the epithelium shows high growth activity is commonly required in theoretical models to reproduce branching patterns [12–14,46]. We examined extracellular signal-regulated kinase (ERK) activity as a hallmark of FGF receptor 2 (FGFR2) activation in embryonic mouse lungs (Fig 1 and S1). Lung explants from a transgenic mouse strain expressing a fluorescence resonance energy transfer (FRET) biosensor for ERK activity were used [38]. To facilitate the laser light for FRET detection in the epithelium, we used a mesenchyme-free epithelium culture supplemented with recombinant FGF10 protein. Fig 1 shows the ERK activity map in the epithelium of the mouse lung at embryonic day 12.5 (E12.5), E13.5, and E14.5, which are the early pseudoglandular stage. It was demonstrated that ERK activity correlates with the cross-sectional curvature of the epithelial explants as *R^2^* = 0.41 in E12.5 and *R^2^* = 0.45 in E13.5, suggesting that FGF10 signaling was enhanced in protruded regions (Fig 1A and 1B). However, poor correlation and activation were observed in the E14.5 epithelium as *R^2^* = 0.0031 (Fig 1C). This result also excluded the possibility that the higher FRET signals at the protruded regions of the earlier explants were not due to differences in detection efficiency. The linear regression analysis revealed a progressive decline in the signal-to-curvature relationship, with a slope of 6.18 at E12.5, 2.93 at E13.5, and 0.12 at E14.5. These results revealed that the signaling response of the epithelium to FGF10 is gradually attenuated with development, indicating the modification of epithelial properties during the developmental process.

We further examined whether the difference in ERK activity reflected its sensitivity to FGF10 concentration (Fig 2). Small explants (<100 µm) of the distal epithelium were grown in Matrigel with various concentrations of FGF10 for 18 h, and the ERK activity of each cyst was measured (panel A in Fig S2). It was demonstrated that FGF10 enhanced ERK activity in a dose-dependent manner, with a correlation of *R^2^* = 0.53 (Fig 2A, black dots). E14.5 cysts showed a poor correlation with *R^2^*= 0.013 (Fig 2A, blue dots), which is consistent with Fig 1C. The slope of regression lines in E13.5 was significantly larger than in E14.5, with *P* = 0.0014, whereas the significant differences in intercepts were rejected, suggesting a higher dose dependency in E13.5 than in E14.5. ERK activation levels vary among cysts for the same FGF10 concentration and the same developmental stage, indicating that epithelial segments may have intrinsic FGF signaling reactivity.

These variations in ERK activity may be due to differences in FGF10-FGFR2 interaction. It was reported that FGF10 stimulates the internalization of FGFR2 as early endosomes [47]. Thus, we visualized FGF10 uptake into cells to observe receptor activation (Fig 2B). Epithelial explants were cultured with fluorescence-labeled FGF10 (FGF10-Alexa488) for 3 h, resulting in the appearance of small fluorescent particles inside the cells (panel B in Fig S2). Immunohistological analysis showed that FGF10 and FGFR2 localization overlapped with the fluorescent signals, suggesting that the signals reflected FGF10-Alexa488 uptake into cells by FGFR2 internalization (Fig S4). Unlike ERK activity, tissue curvature dependency of FGF10 uptake was not detected. The average fluorescent intensity of the explants showed a correlation with the FGF10 dose in E13.5 (*R^2^* = 0.87) and E14.5 (*R^2^* = 0.72), with slopes of 20.7 and 9.7, respectively (Fig 2B). These regression lines differed significantly with *P* = 1.2 × 10^-7^. Collectively, these results suggest that epithelial responsiveness to FGF10 declines with the developmental stages in the lung epithelium, resulting in a difference in the activity gradient depending on the tissue curvature.

### Modeling the curvature-dependent epithelial growth

We considered that epithelial responsiveness to FGF10 may affect the branching pattern and examined this by constructing a mathematical model. The longitudinal section shape of the epithelial duct was described in a 2D space by adopting a model framework, which we tentatively call the particle cell model, in which cells were assumed to be interacting particles [40–44]. We aimed to represent the curvature-dependent growth process of the tissue in the simplest form, in which the FGF10 distribution was omitted.

The cell was assumed to be a uniform circular shape, forming a chain of cells (Fig 3A). Distance-dependent interaction was assumed so that the cells bind to neighboring cells at a fixed distance, as described previously [42,43]. The bending rigidity along the sequence of cells was introduced to express the smooth contours of the tissue as described [42,43]. The interaction strength and bending rigidity are controlled by parameters *k*_wall_ and *k*_bend_, respectively.

Tissue growth was incorporated so that the protruded regions grew faster. Cell migration and division were assumed to depend on local tissue curvature as a phenomenological simplification of epithelial diffusion-limited growth. The local tissue angle *θ_i_* was used instead of calculating the local curvature (Fig 3B). The probability *p_i_* that the *i*-th cell divides in each timestep is as follows:

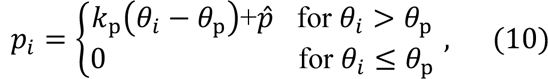

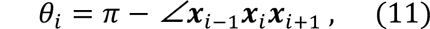

where *k*_p_ is associated with the angle dependence, *θ*_p_ is the minimum angle for the cell division, *p̂* is the basic probability, and ***x****_i_* designates the *i*-th cell position. FGF10 is also known to be involved in the regulation of cell migration [10,19,48]. Here we assumed simple outward movement as follows:

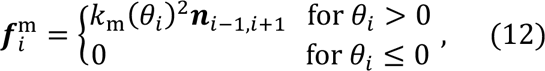

where 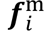 represents the active migration in the normal direction of the tissue, and *k*_m_ is the coefficient.

The cell position changes according to the sum of the calculated forces of the cell-cell interaction, bending rigidity, and migration. The initial shape for the model simulation was assumed to be the tip of the epithelial branch (Fig 3C, left-most image). The cells at the proximal ends were pinned in their initial positions and the distal tip was grown according to curvature-dependent growth, thus depicting the successive tip-splitting deformation (Fig 3C). We further examined how the curvature dependency, denoted by *k*_m_ and *k*_p_, affects branch shape (Fig 3D). It was revealed that larger *k*_m_ and *k*_p_ values resulted in a longer segment length, suggesting that epithelial responsiveness determines the segment length during airway construction. When the curvature dependency was high, extremely restricted tip growth facilitates tip elongation instead of tip splitting. In contrast, in the case of low dependency, broad growth causes tip curvature reduction, causing shape instability. However, the branch width did not show significant differences among the cases with different curvature dependencies, and the entire mechanism of the hierarchy construction could not be explained by the responsiveness change.

### Modeling branch formation with cell shape regulation

To explore the control mechanism of tube size, we focused on the role of cell shape regulation. We have reported that the lung epithelium exhibits apical constriction and facilitates the formation of small buds through Wnt signaling [36]. Inhibition of Wnt signaling has been demonstrated to impede the formation of distal fine branches and induce cystic expansion in mice [31,33]. Drawing from these observations, we considered that apical constriction may facilitate tip splitting to form distal fine branches. To explore this hypothesis, we investigated the impact of apical constriction on branch shape (Fig 4).

We employed the model established in previous studies [36,45], where each cell is represented as a quadrilateral (Fig 4A) to incorporate apical constriction. The model framework is based on the harmonic potential control of each edge length and area, and apical constriction was implemented by applying an attractive force between the pair of apical vertices (Fig 4B). In this study, we extended the model by incorporating curvature-dependent growth. It was confirmed that curvature-dependent growth promoted branch elongation when no apical constriction was assumed (Fig 4C). When apical constriction was assumed, tip splitting was induced (Fig 4D). Apical constriction facilitates tip splitting due to mechanical instability caused by the imbalance between the increase in cell-autonomous curvature due to apical constriction and the decrease in tissue curvature due to growth [15,36,49].

To understand the model dynamics and to explore the mechanisms governing branch width, we depicted small cysts consisting of 13–18 cells and examined the impact of apical constriction (Fig 5). It was demonstrated that apical constriction transformed the cyst from a circular shape to an elongated shape if the cyst contained more than 15 cells; otherwise, the cyst sustained a circular shape (Fig 5A).

Next, we analyzed how the number of cells in the cyst influence the transition from a circular cyst to an elongated shape. The potential energy of the circular state *E_c_* and that of the elongated state *E_e_* were compared (Fig 5B). The longitudinal cyst length, designated by *L*, serves as a defining parameter for the cyst shape. The potential energy of the cyst was assessed based on the regulation of basal length, apical length, and cell area. When it was assumed the basal length and area are sustained by strong regulation, *E_c_*-*E_e_* is derived from the sum of apical lengths as follows:

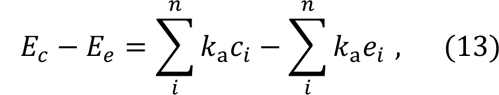

where *n* is the cell number, *k*_a_ represents the strength of apical constriction [36,45], and *c_i_* and *e_i_* are the apical lengths of the *i*-th cell in the circular and elongated cysts, respectively. The sums of the apical lengths were approximated as circular and elliptic circumferences of the cyst lumen as follows:

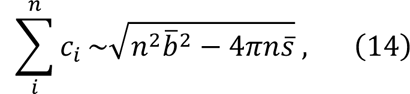

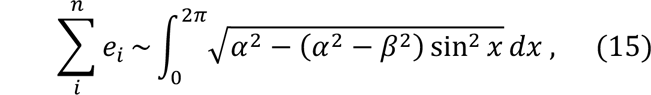

where *b̅* and *s̅* are the basal length and area of the cell, *α* = *L* and *β* are the major and minor radius of an ellipse, respectively. The values of *α* and *β* can be estimated as the function of *b̅*, *s̅*, *n*, and *L* by considering the constant cell area.

Fig 5B shows the numerical profile of *E_c_*-*E_e_* against *L* for *n* ranging from 13 to 18. The condition *n* > 12 is required for *b̅* = *s̅* = 1, as *n*^2^*b̅*^2^ − 4*πns̅* > 0 must be satisfied from Eq (14). The minimum *L* is *b̅n*/*π*, which corresponds to a circular cyst. The maximum of *E_c_*-*E_e_* corresponds to the most stable shape, which is consistent with the model simulation (Fig 5A). The widths of the elongated cysts do not exceed the diameter of the smallest circular cyst, indicating that the same cell shape regulation condition induces spontaneous curvature (Fig 5C). In this regard, we obtained the minimum width among cysts with varying cell numbers and found its decrease with *b̅*/*s̅*, regardless of *s̅* (Fig 5D). This result suggested that cell height determines the spontaneous curvature, irrespective of cell size. Changes in cell size are compensated by alterations in cell number (Fig 5E) and do not affect the spontaneous curvature (Fig 5D).

Fig 5 revealed a correlation between epithelial thickness and branch width, suggesting that the epithelium becomes thinner as development progresses. Supporting this theoretical result, we observed the lung epithelium at E12.5, E13.5, and E14.5, noting a progressive decrease in thickness to 70.3 ± 9.23 µm, 43.0 ± 9.66 µm, and 31.4 ± 4.08 µm, respectively (Fig 6). It should also be noted that E13.5 cysts *in vitro* showed the tendency to be thicker during culture than the E14.5 cysts (Fig S5, black line in panel A and blue solid line in panel C), indicating an intrinsic alteration in thickness regulation as a developmental feature.

### Integrated modeling of curvature-dependent growth and cell shape regulation

In addition to progressive changes in epithelial thickness, curvature-dependent changes in epithelial thickness were also observed. The epithelium was thinner in the high curvature tip region than in the low curvature duct (Fig 6 and 7A), consistent with the detailed 3D analysis of cell shape in E11.5 mouse lung [37], suggesting distinct cell shape regulation regimes for the tip and duct regions. Cells in the concave region were also thickened (asterisks in Fig 7A), implying that the shift from tip-type cells to duct-type cells is occurring during tip spitting event.

**Fig 7.**
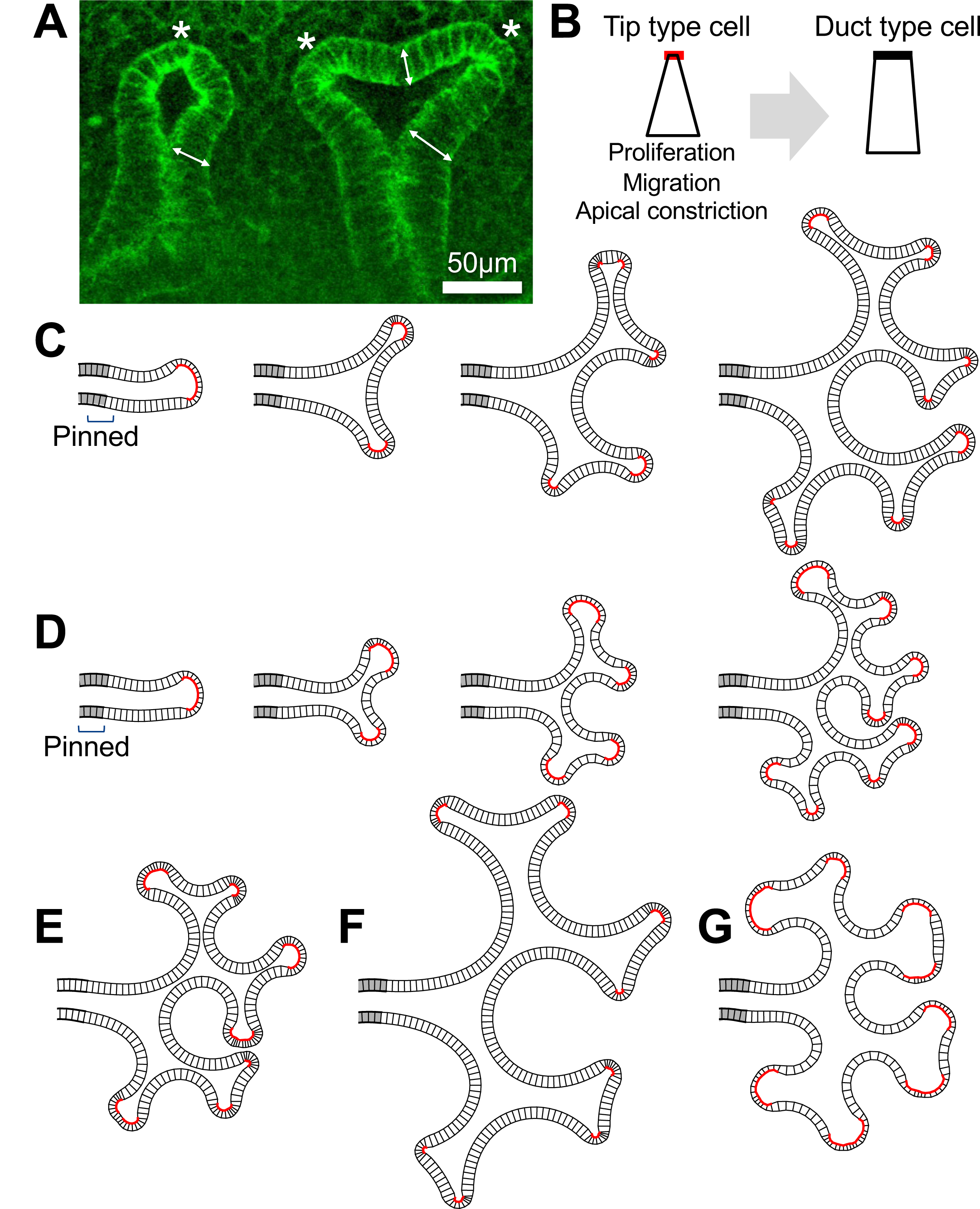
Cell type switching model. **(A)** The growing tip of E14.5 mouse lung. Arrows indicate tall cells at concave regions and asterisks indicate short cells. **(B)** Two cell types in the model. Tip type cells exhibit proliferation, outward migration, and apical constriction (left). Duct type cells are tall and do not show these activities (right). The switch from the tip type to the duct type occurs based on a decrease in local curvature. **(C)** Snapshots of the growing process in the case of thick epithelium with high curvature dependency. The gray color represents cells pinned at their initial positions. The red color on the apical side represents tip type cells. **(D)** Thin epithelium with low curvature dependency. **(E)** Thick epithelium with low curvature dependency, showing the intermediate morphology between (C) and (D). **(F and G)** The cases of low apical constriction activity. Thick epithelium with high curvature dependency (F) and thin epithelium with low curvature dependency (G) are shown.

It has also been shown that canonical Wnt signaling is more intense in the tip region of the lung epithelium [50,51]. We therefore constructed a model to represent that cells in the tip region are highly active in growth and apical constriction.

The model framework is based on the quadrilateral cell model and two types of epithelial cells were assumed: a short tip type exhibiting high growth activity and apical constriction, and a tall duct type that lacks these activities (Fig 7B). The cell type is determined depending on the angle formed by successive vertices, representative of local curvature. The critical angle for switching *φ*_d_ determines the curvature dependency of growth. The growth was described only for the tip type by migration represented by *k*_µ_ and cell division with a constant probability *ρ*. In addition, distinct parameter values for cell shape regulation were assigned to the tip and duct types, which are summarized in S3 Table.

Fig 7C and 7D show that the integration of curvature-dependent growth and cell shape change reproduced the higher-order branching process. Thicker epithelium with a higher curvature dependency resulted in a larger-scale branching structure (Fig 7C), compared to the case of thinner epithelium with a lower curvature dependency (Fig 7D). Epithelial thickness and curvature dependence had an additive impact on branch scale (Fig 7C–E), suggesting that the changes in these epithelial features during development (Fig 1, 2, and 6) contribute to the progressive alteration in airway segment length. It was demonstrated that thin epithelium not only narrowed the branches (Fig 5) but also shortened the branches (Fig 7D and 7E), aligning with the progressive changes in epithelial thickness (Fig 6) and branch morphology [4]. Reduced apical constriction has a more significant effect on the tip shape in thin epithelium compared to thick epithelium (Fig 7F and 7G), consistent with Wnt signaling defects showing severe cystic morphology in the distal airway [33–35].

## Discussion

We explored regulatory mechanisms to generate a hierarchical structure of the lung airways with a specific focus on epithelial properties. Our experimental observations revealed developmental changes in epithelium responsiveness to FGF10. Furthermore, by constructing a mathematical model assuming curvature-dependent growth, we demonstrated that such change leads to variations in branch segment length. Our analysis also highlighted the involvement of cell height regulation in modulating branch width through apical constriction. This study provides a new perspective on how epithelial properties regulate the branching structure of the lung.

Morphogenesis is characterized by cells that are highly active and constantly responding to various regulatory cues, making it an essentially intricate process. Integrating such diverse behaviors into a model can complicate the pursuit of comprehensive understanding. We narrowed down the cellular behaviors of interest based on experimental observations in lung epithelial tissue. The agent-based approach enabled us to explore the implications of curvature-dependent growth and cell shape control in morphogenesis. Our models were based on experimental observations, while the parameters of the model are not experimentally comparable and will need to be validated in the future.

Theoretical studies on lung epithelial morphogenesis have been conducted based on the FGF10 distribution, emphasizing the importance of signaling interaction between the epithelium and mesenchyme [12,14,46]. However, it was later found that the expression pattern of FGF10 in the lung mesenchyme is not required for epithelial branching [52], triggering the hypothesis of an epithelium-autonomous morphogenetic mechanism [15,16,36]. In line with this perspective, our computational models propose that the branching structure of the lung is determined by epithelial properties, FGF10 responsiveness (Fig 3) and tissue thickness (Fig 5). The observation that early epithelia exhibit higher growth activity (Fig 1 and 2), and thicker tissue (Fig 6) might be apparent to biologists studying lung development. Additionally, we found that thinner epithelia are more responsive to FGF10 (panel D in Fig S5), supporting the assumption that the tip type cell is thinner and grows faster in the integrated model (Fig 7B). It was also revealed that FGF10 exposure decreased cyst thickness (panels B and C in Fig S5), implying a potential role of this essential signaling molecule for cell shape regulation.

The regulation of a specific growth activity at the epithelial protrusion remains an unsolved issue (Fig 1). One potentially relevant mechanism is geometry-dependent mechanical patterning [53]. When human embryonic stem cell-derived lung epithelial progenitor cells were cultured inside the 100-µm tube, SOX2 expression was suppressed via canonical Wnt signaling, whereas SOX9 expression was maintained [53]. The tissue curvature-dependent regulation of SOX2 and SOX9 expression, which is regulated only downstream of the Wnt/b-catenin pathway, is consistent with the assumption of curvature-dependent growth and apical contraction in our model (Fig 7). Clarifying how the cell shape and growth activity are regulated and how signaling pathways such as FGF and Wnt are involved is necessary to gain insight into the regulatory mechanisms of the global structure of the lung.

## Author Contributions

H.T.-I. designed research, performed experimental and theoretical research, constructed models, and wrote the manuscript; H.T. executed experiments and analyzed data; K.F. and T.M. supervised research. All authors have read and approved the published manuscript.

## Acknowledgment

We thank Prof. Michiyuki Matsuda (Kyoto University) for the use of Eisuke transgenic mice. We thank Ms. Yoshimi Yamaguchi for her technical assistance with the FGF10 labeling experiment. This work was supported by JSPS KAKENHI Grant Number 18K06260 (H.T.-I.) and 15KT0018 (T.M).

