## Supporting Information for "Exploiting mechanisms for hierarchical branching structure of lung airway"

### FIG S1

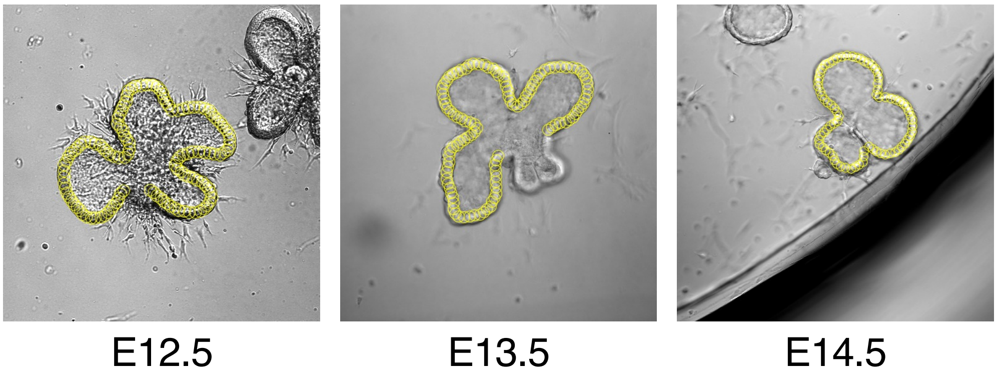


**Fig S1. ROIs used to obtain the ERK activity map in Fig 1.** ROIs are indicated by yellow circles on the bright-field images. 2D tissue curvature and ERK activity were obtained from a series of ROI clockwise along the epithelial cross-section.

### FIG S2

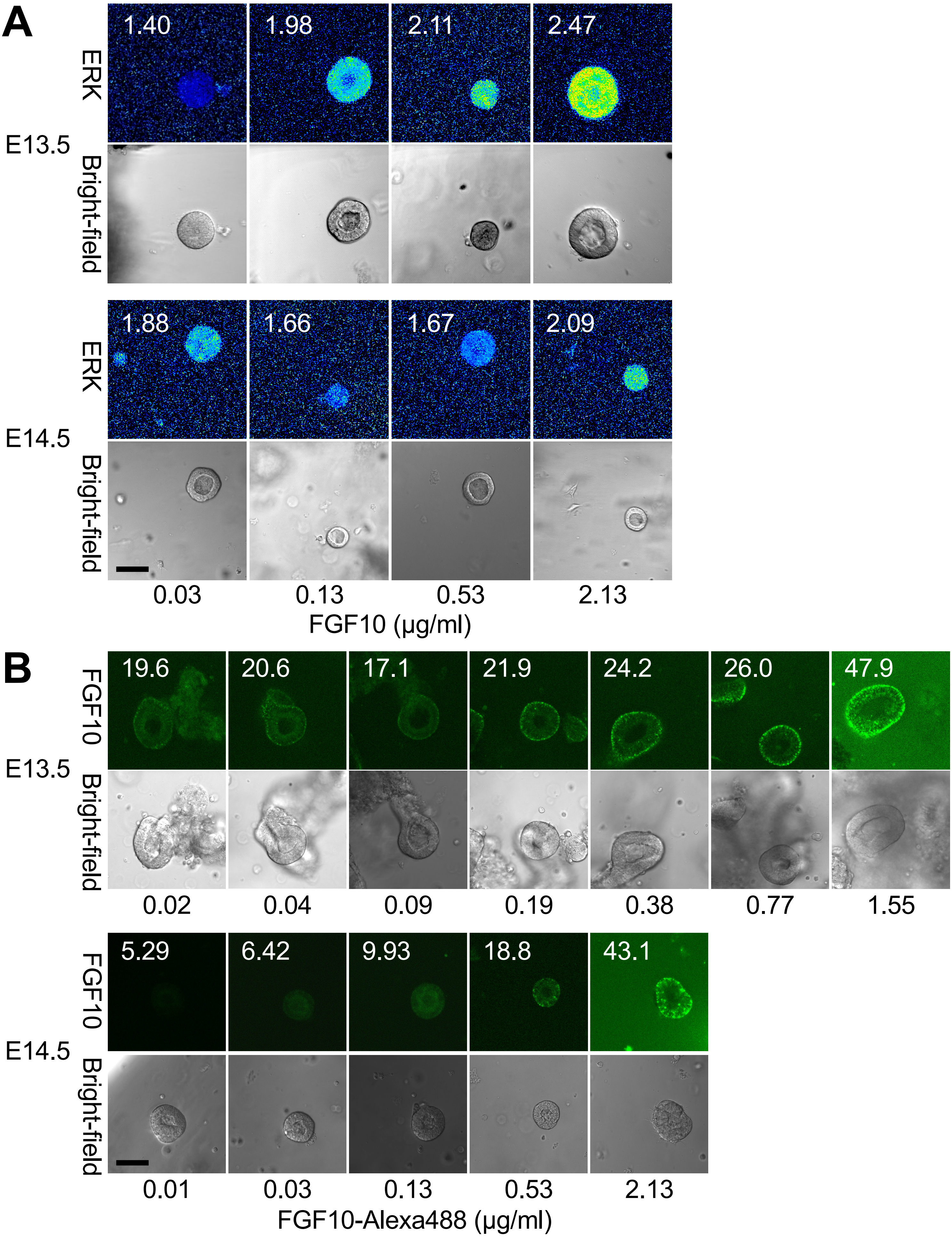

A
E13.5
ERK
Bright-field
1.40
1.98
2.11
2.47
E14.5
ERK
Bright-field
1.88
1.66
1.67
2.09
0.03
0.13
0.53
2.13
FGF10 (µg/ml)
B
E13.5
FGF10
Bright-field
19.6
20.6
17.1
21.9
24.2
26.0
47.9
0.02
0.04
0.09
0.19
0.38
0.77
1.55
E14.5
FGF10
Bright-field
5.29
6.42
9.93
18.8
43.1
0.01
0.03
0.13
0.53
2.13
FGF10-Alexa488 (µg/ml)

**Fig S2. Representative images in the ERK activity measurement in E13.5 and E14.5 lung epithelial explants in Fig 2. (A)** The heat maps of the ERK activity (upper panels) and bright field images (lower panels) for E13.5 and E14.5 cysts. The ERK activity quantified for the cyst is indicated on each panel. The epithelial thickness seems different between in E13.5 and E14.5, and the statistical analysis was demonstrated in Fig S3. **(B)** The fluorescent signals of FGF10-Alexa488 (upper panels) and bright field images (lower panels) for E13.5 and E14.5 explants. High backgrounds of the fluorescent images were caused by the high FGF10-Alexa488 concentration in the Matrigel and the medium. The mean fluorescent intensity is indicated on each panel. Scale bar: 50 µm.

**FIG S3**

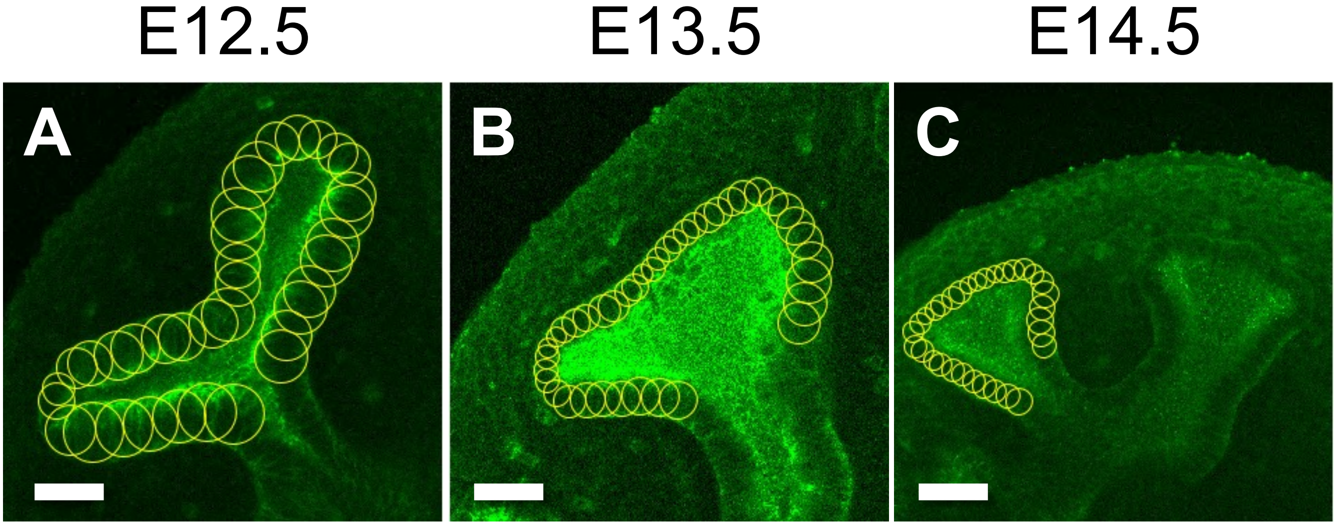


**Fig S3. ROIs used to measure the epithelial thickness in Fig 6.** ROIs for thickness measurement. Small circles were fitted to the local thickness of the epithelium. The mean and standard deviation of the circle diameters were calculated to represent the thickness.

FIG S4

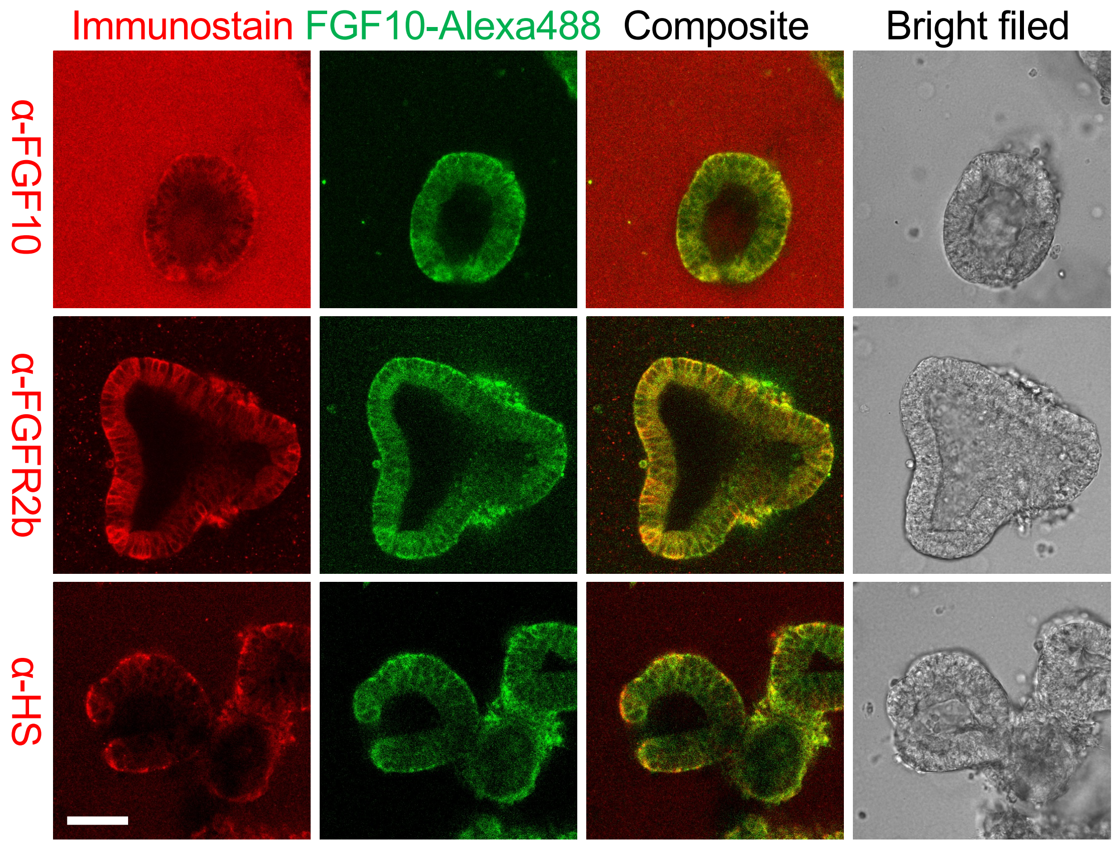


**Fig S4. Immunohistological analysis validating the FGF10-Alexa488 uptake experiment as the indication of the FGF10-FGFR2 internalization.** The E13.5 epithelial cyst was cultured with the FGF10-Alexa488 and subsequently applied to the immunostaining of FGF10, FGFR2, and heparan sulfate. The samples were fixed with 2% paraformaldehyde in PBS for 5 min on ice. We used anti-FGF10 antibody (H-121, 1:20; Santa Cruz), anti-Bek (C-17, 1:100; Santa Cruz) for FGFR2 detection, and anti-heparan sulfate (10E4, 1:100; Seikagaku). The second antibodies were Alexa Fluor 568-conjugated goat anti-rabbit IgG and goat anti-mouse IgG (1:200; Thermo Fisher). The Alexa488 signals in cells was diffused but remained after the staining. The result of FGF10 immunostaining coincided with the FGF10-Alexa488 signal, except for that Matrigel containing FGF10-Alexa488 strongly reacted to anti-FGF10 antibody (upper panels). It was also found that FGFR2 mostly colocalized to the FGF10-Alexa488 signals (middle panels). Heparan sulfate is known to be localized to the cell surface and to support the FGF10-FGFR2 binding (lower panels). Scale bar: 50 μm.

FIG S5

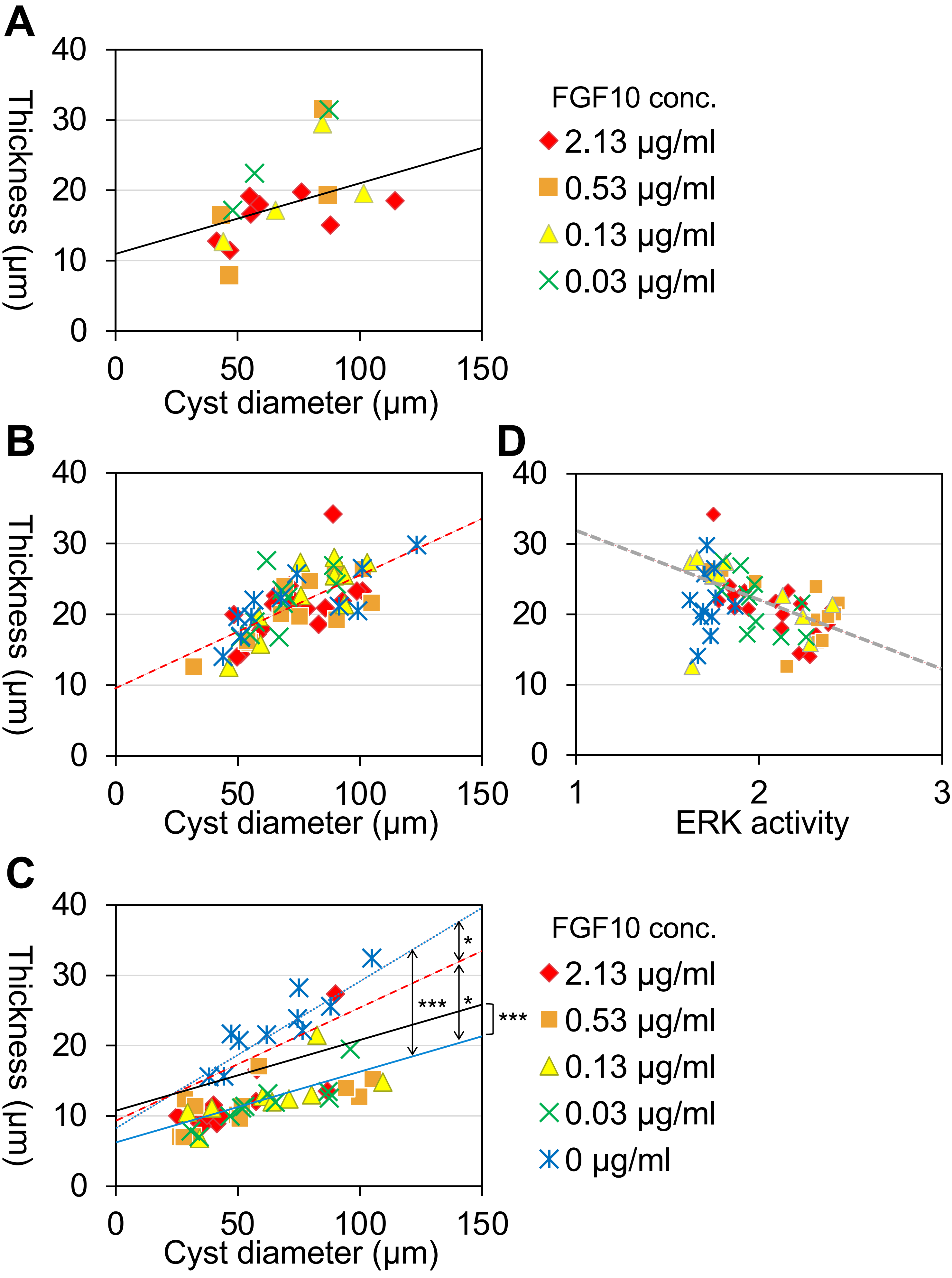
A
B
C
D
Thickness (μm)
Cyst diameter (μm)
ERK activity
FGF10 conc.
2.13 μg/ml
0.53 μg/ml
0.13 μg/ml
0.03 μg/ml
0 μg/ml
*
***
0
10
20
30
40
50
100
150
1
2
3

**Fig S5. Shape changes and ERK activity of lung epithelial cysts.** The cysts from E13.5 (A) and E14.5 (B, C, and D) mice were cultured in the FGF10-supplimented Matrigel for 18 h (A and C) or 3 h (B and D), following the procedures outlined in in Fig 2 and S2. The series for E13.5 and E14.5 are presented to the right of (A) and (D), respectively. For E13.5, morphometric analysis could not be conducted on samples cultured for 3 h with FGF and for 18 h without FGF, as cyst formation in these cases showed minimal progress. Cyst thickness was quantified by estimating outer and inner diameters in cross-sections using Fiji (NIH), calculated as the difference between them. The statistical analysis was performed by using R. The significant difference among the linear regression lines was tested by the analysis of covariance (ANOVA). **(A)** E13.5 cysts cultured for 18 h. The thickness-to-diameter correlations in four different doses of FGF10 showed no significant difference. The linear regression for the pooled data is $y = 0.10x + 11.39$ with $R^2 = 0.24$ (black line), suggesting that epithelia tend to be thicker in larger cysts. Small explants may have been obtained by tearing off the tip of the epithelium where the epithelium is thin, while large explants may have been from a large area containing a thicker duct. **(B)** E14.5 cysts cultured for 3 h. The correlations in five different doses showed no significant difference. The linear regression for the pooled data is $y = 0.16x + 9.48$ with $R^2 = 0.48$ (red dashed line), confirming the same tendency in (A). **(C)** E14.5 cysts cultured for 18 h. The correlation in five different doses showed no significant difference except for the samples without FGF10 ($P < 0.001$), suggesting that FGF10 exposure flattened the epithelium. The linear regression for the pooled data of the FGF10 exposed cases is $y = 0.10x + 9.48$ with $R^2 = 0.48$ (blue line), and for the case without FGF10 $y = 0.21x + 8.24$ with $R^2 = 0.79$ (blue dotted line). Both correlations were significantly different from the case of 3-h culture in (B) ($P < 0.05$), confirming that FGF10 exposure progressively flattened the epithelium, whereas prolonged culture without FGF10 thickened the epithelium. These correlations are also significantly different from the case of E13.5 cysts in (A), supporting the difference of the epithelial properties along the developmental process. The slopes in (A) and pooled data in (B) did not differ, whereas their intercepts significantly differ by 5.14 ($P < 0.001$), which agrees with that epithelium in the earlier stage is thicker. **(D)** Thickness was plotted against ERK activity for the corresponding sample in (B). The linear regression for the pooled data of the FGF10 exposed cases is $y = -10.08x + 42.10$ with $R^2 = 0.28$ (gray dotted line), indicating that thinner epithelia exhibit higher activity. No significant correlation was found for the samples without FGF10.

### Table S1

| Parameter and description | | Fig 3C | Fig 3D |
|---|---|---|---|
| $k_\mathrm{p}$ | Angle dependence of cell division | 0.008 | * |
| $\theta_\mathrm{p}$ | Minimum angle for cell division | 0.2 | 0.2 |
| $\hat{p}$ | Basic probability | 0.001 | 0.001 |
| $k_\mathrm{m}$ | Angle dependence of active migration | 25 | * |
| $k_\mathrm{wall}$ | Interaction between adjacent cells [42] | 60 | 60 |
| $k_\mathrm{bend}$ | Bending rigidity along the cell sequence [42] | 15 | 15 |
| $\gamma$ | Friction coefficient for cell displacement | 15 | 15 |

*: Values are indicated in the figure.

### Table S2

| Parameter and description | | Fig 4C | Fig 4D | Fig 5A |
|---|---|---|---|---|
| $k_{\mathrm{q}}$ | Dependency of cell division on the angle of the basal contour | 0.9 | 0.9 | 0 |
| $\varphi_{\mathrm{q}}$ | Minimum angle for cell division | 0.5 | 0.5 | 0 |
| $\hat{q}$ | Basic probability for cell division | 0.0001 | 0.0001 | 0 |
| $k_{\mathrm{w}}$ | Dependency of active migration on the angle of the basal contour | 1 | 1 | 0 |
| $\varphi_{\mathrm{w}}$ | Minimum angle for the active migration | 0.7 | 0.7 | 0 |
| $\bar{a}$ | Optimal apical length of the cell [36] | 0 | 0 | 0 |
| $\bar{b}$ | Optimal basal length of the cell [36] | 1 | 1 | 1 |
| $\bar{s}$ | Optimal area of the cell | 1 | 1 | 1 |
| $k_{\mathrm{a}}$ | Apical length regulatory coefficient [36] | 0 | 1 | ** |
| $k_{\mathrm{b}}$ | Basal length regulatory coefficient [36] | 300 | 300 | 300 |
| $k_{\mathrm{s}}$ | Area regulatory coefficient | 3 | 3 | 3 |
| $k_c$ | Lateral length regulatory coefficient [36] | 0 | 0 | 0 |
| $k_{\mathrm{ab}}$ | Cell shape symmetry coefficient [36] | 10 | 10 | 5 |
| $k_{\mathrm{bend}}$ | Bending rigidity of the basal contour [36] | 0.2 | 0.2 | 2 |
| $\gamma$ | Friction coefficient | 20 | 20 | 20 |

**: The value is 0 during relaxation without apical constriction and 3 for applying apical constriction.

### Table S3

| Parameter and description | | Fig 7C | Fig 7D | Fig 7E | Fig 7F | Fig 7G |
|---|---|---|---|---|---|---|
| $\varphi_{\mathrm{d}}$ | Critical angle for cell type switching | -0.45 | -0.3 | -0.3 | -0.45 | -0.3 |
| $k_{\mu}$ | Active migration coefficient for tip type | 0.8 | 0.6 | 0.6 | 0.8 | 0.6 |
| $\rho$ | Cell division probability for tip type | 0.00025 | 0.0002 | 0.0002 | 0.00025 | 0.0002 |
| $\bar{a}_{\mathrm{t}}$ | Optimal apical length [36] for tip type | 0 | 0 | 0 | 0 | 0 |
| $\bar{a}_{\mathrm{d}}$ | Optimal apical length [36] for duct type | 1.1 | 1.4 | 1.1 | 1.1 | 1.4 |
| $\bar{b}$ | Optimal basal length [36] | 1.1 | 1.4 | 1.1 | 1.1 | 1.4 |
| $\bar{s}_{\mathrm{t}}$ | Optimal cell area for tip type | 1 | 1 | 1 | 1 | 1 |
| $\bar{s}_{\mathrm{d}}$ | Optimal cell area for duct type | 2.7 | 2.7 | 2.7 | 2.7 | 2.7 |
| $k_{\mathrm{at}}$ | Apical length regulatory coefficient [36] for tip type | 70 | 70 | 70 | 35 | 35 |
| $k_{\mathrm{ad}}$ | Apical length regulatory coefficient [36] for duct | 40 | 40 | 40 | 40 | 40 |
| $k_{\mathrm{b}}$ | Basal length regulatory coefficient [36] | 300 | 300 | 300 | 300 | 300 |
| $k_{\mathrm{s}}$ | Area regulatory coefficient | 50 | 50 | 50 | 50 | 50 |
| $k_{\mathrm{c}}$ | Lateral length regulatory coefficient [36] | 0 | 0 | 0 | 0 | 0 |
| $k_{\mathrm{ab}}$ | Cell shape symmetry coefficient [36] | 200 | 200 | 200 | 200 | 200 |
| $k_{\mathrm{bend}}$ | Bending rigidity of the basal side | 30 | 30 | 30 | 30 | 30 |
| $\gamma$ | Friction coefficient | 20 | 20 | 20 | 20 | 20 |
